# Individualized precision targeting of dorsal attention and default mode networks with rTMS in traumatic brain injury-associated depression

**DOI:** 10.1101/2021.03.13.435127

**Authors:** Shan H. Siddiqi, Sridhar Kandala, Carl D. Hacker, Nicholas T. Trapp, Eric C. Leuthardt, Alexandre R. Carter, David L. Brody

**Affiliations:** Department of Psychiatry, Washington University School of Medicine, 660 S Euclid Ave, St. Louis, MO 63110; Berenson-Allen Center for Noninvasive Brain Stimulation, Harvard Medical School, 330 Brookline Ave, Boston, MA 02215; Center for Neuroscience and Regenerative Medicine, Uniformed Services University of the Health Sciences, 4301 Jones Bridge Rd, Bethesda, MD 20814; Department of Neurosurgery, Washington University School of Medicine, 660 S Euclid Ave, St. Louis, MO 63110; Department of Psychiatry, University of Iowa Carver College of Medicine, 500 Newton Rd, Iowa City, IA 52246; Department of Neurology, Washington University School of Medicine, 660 S Euclid Ave, St. Louis, MO 63110

**Keywords:** transcranial magnetic stimulation, traumatic brain injury, depression, individualized parcellation, resting state network, neuronavigation, fMRI

## Abstract

**Background:** At the group level, antidepressant efficacy of rTMS targets is inversely related to their normative connectivity with subgenual anterior cingulate cortex (sgACC). Individualized connectivity may yield better targets, particularly in patients with neuropsychiatric disorders who may have aberrant connectivity. However, sgACC connectivity shows poor test-retest reliability at the individual level. Individualized resting-state network mapping (RSNM) can reliably map inter-individual variability in brain network organization.

**Objective:** To identify individualized RSNM-based rTMS targets that reliably target the sgACC connectivity profile.

**Methods:** We used RSNM to identify network-based rTMS targets in 10 healthy controls and 13 individuals with traumatic brain injury-associated depression (TBI-D). These “RSNM targets” were compared with consensus structural targets and targets based on individualized anti-correlation with a group-mean-derived sgACC region (“anti-group-mean sgACC targets”). The TBI-D cohort was randomized to receive active (n=9) or sham (n=4) rTMS to RSNM targets.

**Results:** The group-mean sgACC connectivity profile was reliably estimated by individualized correlation with default mode network (DMN) and anti-correlation with dorsal attention network (DAN). Individualized RSNM targets were then identified based on DAN anti-correlation and DMN correlation. Counterintuitively, anti-correlation with the group-mean sgACC connectivity profile was stronger and more reliable for RSNM-derived targets than for “anti-group-mean sgACC targets”. Improvement in depression after RSNM-targeted rTMS was predicted by target anti-correlation with the portions of sgACC. Active treatment led to increased connectivity within and between several relevant regions.

**Conclusions:** RSNM may enable reliable individualized rTMS targeting, although further research is needed to determine whether this personalized approach can improve clinical outcomes.

## 1. Introduction

The antidepressant efficacy of repetitive transcranial magnetic stimulation (rTMS) may be related to the connectivity of the stimulation target^1^. Most commonly, scalp measurements or structural MRI are used to identify a target in the dorsolateral prefrontal cortex (DLPFC)^2^. Recent studies have attempted to identify rTMS targets based on functional connectivity (FC) with “seed” regions deeper in the brain^3^. At the group level, antidepressant efficacy of rTMS is related to normative anti-correlation between the stimulation target and subgenual anterior cingulate cortex (sgACC), suggesting that treatment may be suppressing activity in sgACC and the limbic system^4-7^. To improve upon these group-level targets, individualized connectivity measurements have also been used to identify patient-specific stimulation sites^8-10^. These targets may be superior to normative “anti-group mean sgACC” targets^6,7,11^.

However, such targeting approaches are limited by the fact that sgACC connectivity is unreliable at the individual level^8,10,12^. Reliability assessments have shown weak test-retest correlation between sgACC connectivity maps (spatial r<0.5)^12^ and marked variability in DLPFC targets identified based on sgACC connectivity (mean test-retest variability of 25mm)^10^. Targets can be identified more reliably based on connectivity with the “network” of regions most correlated with the sgACC at the group level^8^. This network may be personalized using individualized resting-state network mapping (RSNM), which can reliably map brain networks based on resting-state functional MRI (rsfMRI)^13-17^. RSNM has been successfully used for neurosurgical pre-operative mapping^18^ and has recently been evaluated as a method for mapping prefrontal topography to identify rTMS targets^19^.

RSNM enables precise individualized mapping of the DMN^20^, which is highly correlated with sgACC and the limbic system^21^. Individualized DMN mapping may thus serve as a more reliable proxy for sgACC connectivity. DMN is strongly anti-correlated with dorsal attention network (DAN)^22-24^, so individualized DAN mapping may yield a TMS target that is reliably anti-correlated to sgACC. Indeed, RSNM studies have found that DAN usually includes a node in the DLPFC, but the precise location of this node varies greatly between individuals^25,26^. Reliable rTMS targets have been identified at this node^19^. Stimulation of this node led to changes in sgACC connectivity with the DAN stimulation sites and with the DMN^19^.

Of note, much of the existing knowledge about individualized brain mapping has been based on studies of healthy individuals. It remains unclear whether this generalizes to patients with neuropsychiatric illnesses and brain injuries. Inter-individual variability may be particularly prominent in traumatic brain injury-associated depression (TBI-D), which is associated with altered FC in the DLPFC, sgACC, DAN, and DMN^27-30^. This raises additional questions regarding the appropriateness of group-mean rTMS targets or individualized targets derived from seed-based connectivity.

As a first step to addressing these questions, we explored the differences between potential target sites generated using individualized RSNM (RSNM-based targets), standard anatomical methods (structural targets), and the point of maximal anti-correlation with the group-mean location of the sgACC (anti-group mean sgACC targets). We also explored the connectivity changes induced by stimulation of RSNM-based targets in a recent pilot clinical trial^19^. We hypothesized that RSNM-based targets would approximate the sgACC connectivity profile more reliably than a group-based sgACC seed, which has previously been proposed for rsfMRI-based rTMS targeting^6,8,10^. After stimulation of these RSNM-based targets, we hypothesized that connectivity changes would be observed in the targeted networks, that these connectivity changes would covary with antidepressant response, and that antidepressant response would be predicted by baseline sgACC connectivity to the stimulation site.

## 2. Methods

Full methodological details are presented in the supplement.

### 2.1 Standard protocol approvals and participants

Data were collected as part of a pilot trial of rTMS for TBI-D^19^. Methods and hypotheses were pre-registered with the Open Science Foundation (osf.io/vjddq)^31^ and ClinicalTrials.gov (NCT02980484).

15 subjects (11 males, ages 19-64) were recruited if they scored at least 10 on the Montgomery-Asberg Depression Rating Scale (MADRS) and had a history of at least one concussive or moderate TBI. This analysis was limited to 13 subjects (10 males) who completed both the pre-treatment and post-treatment scan sessions.

10 healthy control subjects (3 males, ages 22-35) with no history of neuropsychiatric disease were chosen randomly from the Human Connectome Project (HCP) database^32^. Only 10 subjects were chosen in order to approximately match the number of TBI-D patients and to confirm that utility of individualized RSNM can be demonstrated with small sample sizes; clinical practicality of personalized medicine approaches may be questionable if large sample sizes are required to demonstrate their utility.

### 2.2 MRI acquisition and pre-processing

For TBI-D subjects, a 3T Siemens Magnetom Prisma magnetic resonance scanner was used to acquire 16.5 minutes of resting-state blood oxygen-level dependent (BOLD) data in three runs. For HCP subjects, a 3T Siemens Connectome Skyra was used to acquire 58 minutes of resting-state BOLD data in four runs. Preprocessing was conducted using in-house scripts described in Power et al., 2014^33^. For each subject, BOLD time courses were used to construct seven individual-level RSN maps via a multilayer perceptron (MLP)-based machine learning classifier as described in Hacker et al., 2013^15^. To create individualized regions of interest for further analyses, a winner-take-all map was created by assigning each voxel to the network with maximum likelihood of membership. Further details are described in the supplement.

### 2.2 Confirmation of candidate network targets

#### 2.3.1 Normative connectivity data

Data from the HCP 800-subject release^32^ were used to construct normative maps of resting-state functional connectivity with the sgACC (figure 1a), as described in Weigand et al., 2018^5^. We hypothesized that the individualized map of the DAN would be most similar to the normative map of sgACC anti-correlations. We also hypothesized that the individualized map of the DMN would be most similar to the normative map of sgACC positive correlations. To test these hypotheses, the group-based map was compared with individualized RSN maps for each subject (figure 1b). All subsequent analyses were conducted using subject-specific connectivity data rather than group connectome data.

**Figure 1.**
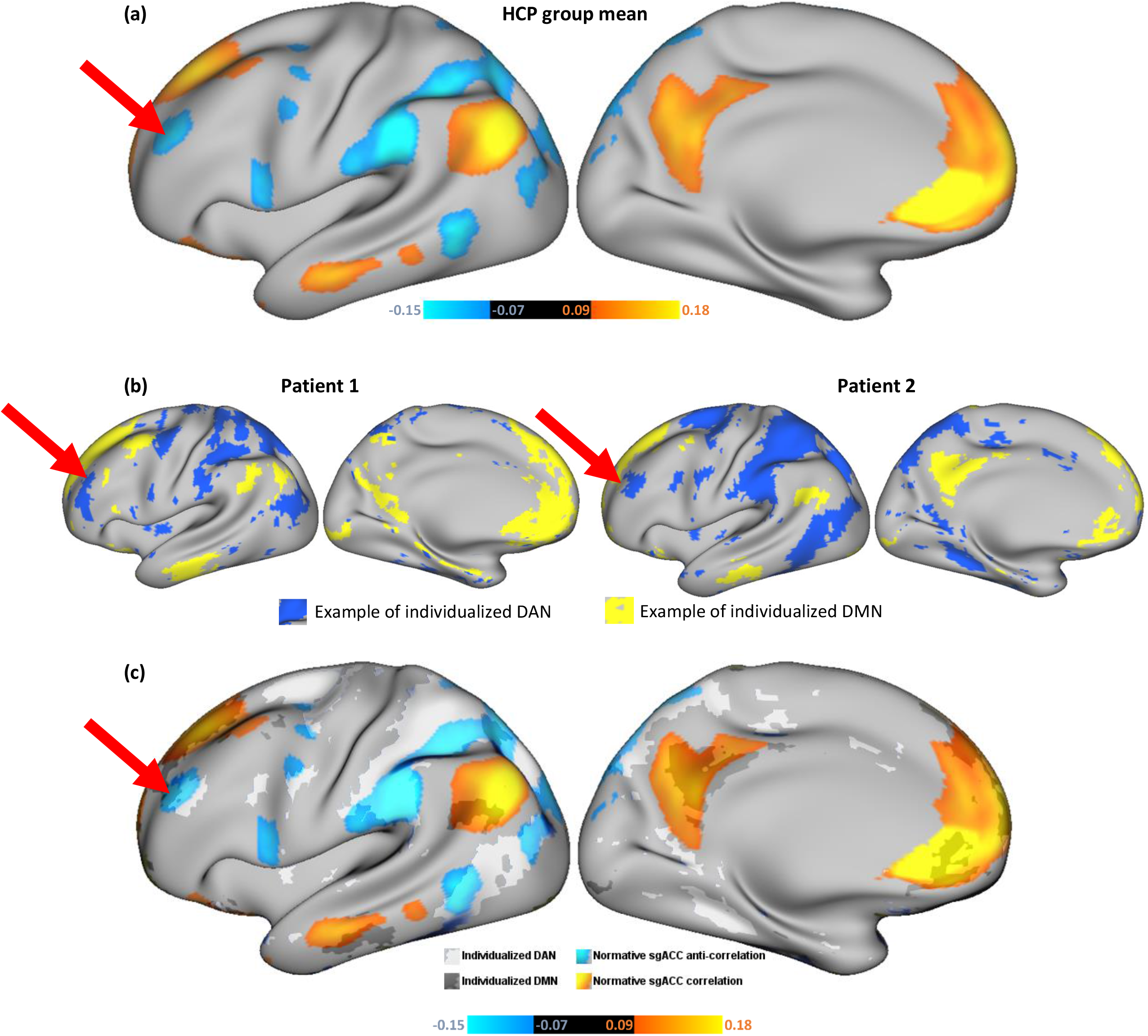
**(a)** Normative map of sgACC functional connectivity, indicating strongest areas of correlation (orange-yellow), and anti-correlation (blue). Strong normative sgACC anti-correlation is prominent at a DLPFC site (red arrow) which has previously been shown to be an effective rTMS target in major depression^4,5^. **(b)** Individualized winner-take-all maps of DAN (blue) and DMN (yellow) for two representative example subjects. Red arrows depict the group-mean stimulation site, which shows differing spatial relationships with DAN in the two patients **(c)** Example of overlap between individualized RSNM DAN/DMN maps (from patient 2) and normative sgACC seed map.

#### 2.3.2 Comparison of individualized RSN maps to sgACC seed maps

For each subject, the seven individualized RSN maps were compared with the normative sgACC seed map. The sgACC seed map was masked with each individual RSN map to identify its overlap with that network (figure 1c). This yielded a map of normative sgACC connectivity values at each voxel within each RSN. Using this map, the mean normative sgACC connectivity value of overlapping voxels was calculated for each RSN and each subject. This yielded a quantitative metric of the degree of overlap between the continuous normative sgACC seed map and each binary RSN map. For each RSN, this value was compared with DAN and DMN by calculating Fisher’s least significant difference via one-way ANOVA.

### 2.4 rTMS target selection and comparison

Specific analytical procedures/tools are described in the supplement.

#### 2.4.1 Target selection

Three approaches were used to identify potential rTMS targets:

1. Individualized RSNM-based targeting – The individualized DMN map was subtracted from the individualized DAN map for each subject. The peak DLPFC cluster was identified in this map following the methods described in Siddiqi et al., 2019 (Fig. S1)^34^.
2. Structural MRI-based targeting – Targets were chosen at DLPFC coordinates (±38, 44, 26), which have been used for targeting at the world’s current largest neuronavigated rTMS clinic^35^.
3. Individualized anti-group mean sgACC target – this method relies on an individual subject’s anti-correlation with group-mean sgACC coordinates, as described in Fox et al., 2013^8^. Similar approaches have been implemented in at least two recent prospective studies^9,10^.

#### 2.4.2 Comparison of resting state functional connectivity of the potential targets

For each potential stimulation site, resting-state functional connectivity was calculated with a consensus group-mean definition of DAN and DMN^36^. To confirm that effects were not driven by autocorrelation between the RSNM-based DAN/DMN parcels and the consensus DAN/DMN parcels, connectivity was also calculated with the normative sgACC seed map. If effects were driven by such autocorrelation, then the sgACC seed map would be most anti-correlated with the anti-group mean sgACC targets.

Potential target correlations with the DAN, DMN, and the sgACC seed map were compared between the different targeting methods across all subjects via within-subjects two-way ANOVA. Results from the two groups of subjects (TBI-D and healthy controls) were not compared with one another due to potential influence of methodological variability and demographic differences.

#### 2.4.3 Comparison of spatial locations of the potential targets

For each subject, the three potential targets were also compared in terms of spatial distance between one another. The mean distances of RSNM targets and anti-group mean sgACC targets from the structural target were compared using paired t-tests. Inter-individual variances for RSNM targets and anti-group mean sgACC targets were determined using F-tests based on the distance of each target from the mean of all coordinates generated by that method.

### 2.5 rTMS treatment

To explore the effects of stimulating our proposed targets, TBI-D subjects were randomized to receive 20 daily sessions of active or sham rTMS using the RSNM targets. The clinical treatment protocol and results are described in detail in Siddiqi *et al*, 2019^19^. Briefly, clinical treatment included 4000 pulses of high-frequency (10 Hz) left-sided stimulation, followed by 1000 pulses of low-frequency (1 Hz) right-sided stimulation. Using a Brainsight neuronavigation device, target coordinates were plotted on a surface reconstruction of each subject’s brain. No stimulation at other targets was performed in the current study.

### 2.6 Treatment-induced changes

Detailed analysis parameters are described in the supplement.

#### 2.6.1 Target stability over time

Nine TBI-D subjects were randomized to active treatment and four were randomized to sham. For each of the three targeting methods, connectivity with the normative sgACC seed map was calculated for pre-treatment and post-treatment scans. Two-way ANOVA was used to compare the three targeting methods in terms of difference in connectivity between the two time points. Again, it should be noted that only individualized RSNM-based targeting was performed.

Euclidean distances between pre-treatment and post-treatment targets were calculated for RSNM and anti-group mean sgACC targets in order to assess the stability of target location. Due to non-normal distribution of these distances, a Wilcoxon matched-pairs signed-rank test was used to compare the two targeting methods in terms of stability over time.

#### 2.6.2 Treatment-induced connectivity changes

In an exploratory analysis, active and sham groups were compared in terms of treatment-induced change in connectivity. Connectivity was calculated between five *a priori* ROIs defined in the original clinical trial, including left/right stimulation sites, DAN, DMN, and sgACC. This analysis was conducted using covariance rather than correlation, since covariance is less sensitive to the potential influence of changing amplitudes of BOLD fluctuations between different time points. In two exploratory analyses, ROI-ROI connectivity was calculated with each of the 17 Yeo networks and voxel-wise connectivity was calculated with the whole brain. Active and sham groups were compared using a general linear model (GLM) with group assignment as the primary predictor, post-treatment connectivity as the outcome, and pre-treatment connectivity as a covariate. Except where required for voxel-wise multiple comparisons correction, statistical hypothesis testing was not conducted for active-sham comparisons because the trial did not reach its original target sample size^19^.

#### 2.6.3 Prediction of antidepressant response

To examine connectivity-based predictors of response in the active treatment group, whole-brain connectivity of each stimulation site was compared with antidepressant response. For each voxel, a least squares regression model was constructed using baseline target-voxel connectivity and baseline MADRS as predictors of post-treatment MADRS. Because antidepressant response could not be assumed to be normally distributed in this small sample, all data were rank-transformed, which is consistent with prior methods described in Weigand *et al*, 2018^5^.

## 3. Results

### 3.1 Confirmation of candidate network targets

In both groups, the positive correlations in the normative sgACC seed map (figure 1a, yellow/orange regions) showed stronger overlap with the individual DMN map than with any other individualized network map (figure 2a). The anti-correlations in the sgACC seed map (Fig. 1a, blue regions) showed stronger overlap with the individualized DAN map than with any other network map (figure 2a). In comparison with the other individualized networks in the TBI-D group, DAN showed significantly stronger overlap with the negative component of the sgACC seed map, while DMN showed significantly stronger overlap with the positive component of the sgACC seed map (figure 2b). The same trend was evident in the HCP group, except that the DAN-ventral attention network (VAN) difference and the DAN-frontoparietal control network (FPC) difference did not reach significance (figure 2b). Overall, DAN anti-correlation and DMN correlation provided the best individualized approximation of the sgACC seed map.

**Figure 2:**
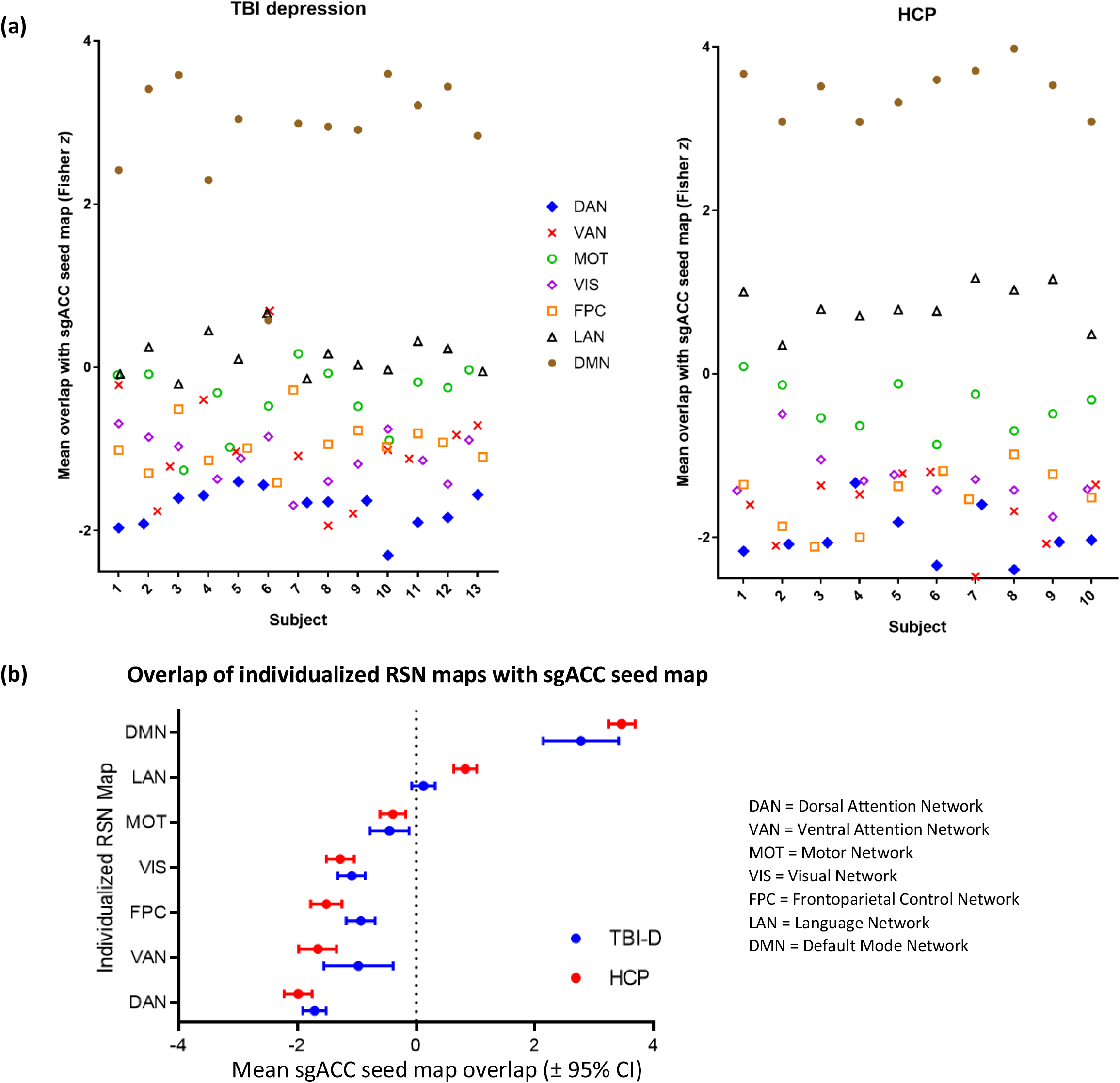
**(a) Individualized similarity between each RSN and the normative sgACC seed map at baseline**. The group-mean sgACC seed map positive correlations overlapped more with DMN and the anti-correlations overlapped more with DAN than any other individualized RSN map for the majority of individual subjects. **(b) Mean similarity between individualized RSNs and the normative sgACC seed map**. DMN was the only network showing strong overlap with the positive correlations in the sgACC seed map. Several RSNs showed notable overlap with the anti-correlations in the sgACC seed map.

### 3.2 Evaluation of expected stimulation profile for each potential target

Nearly all potential targets showed positive correlation with the DAN, negative correlation with the DMN, and negative correlations with the group mean sgACC seed map (figure 3a). Within-subjects two-way ANOVA revealed a significant effect of potential targeting method on left- and right-sided target connectivity with each of these regions in each of the two datasets (table S1). The magnitude of these differences was similar but not identical between the two datasets (figure 3b and table S1). In comparison with each of the other two targets, the RSNM-based target showed stronger DAN correlation and DMN anti-correlation in 11/13 TBI-D patients and 8/10 healthy controls (p=0.002, single-proportion z-test with expected proportion of 50%). Overall, RSNM-based targets appeared to provide a better individualized approximation of the desired networks than the other two potential targeting methods.

**Figure 3:**
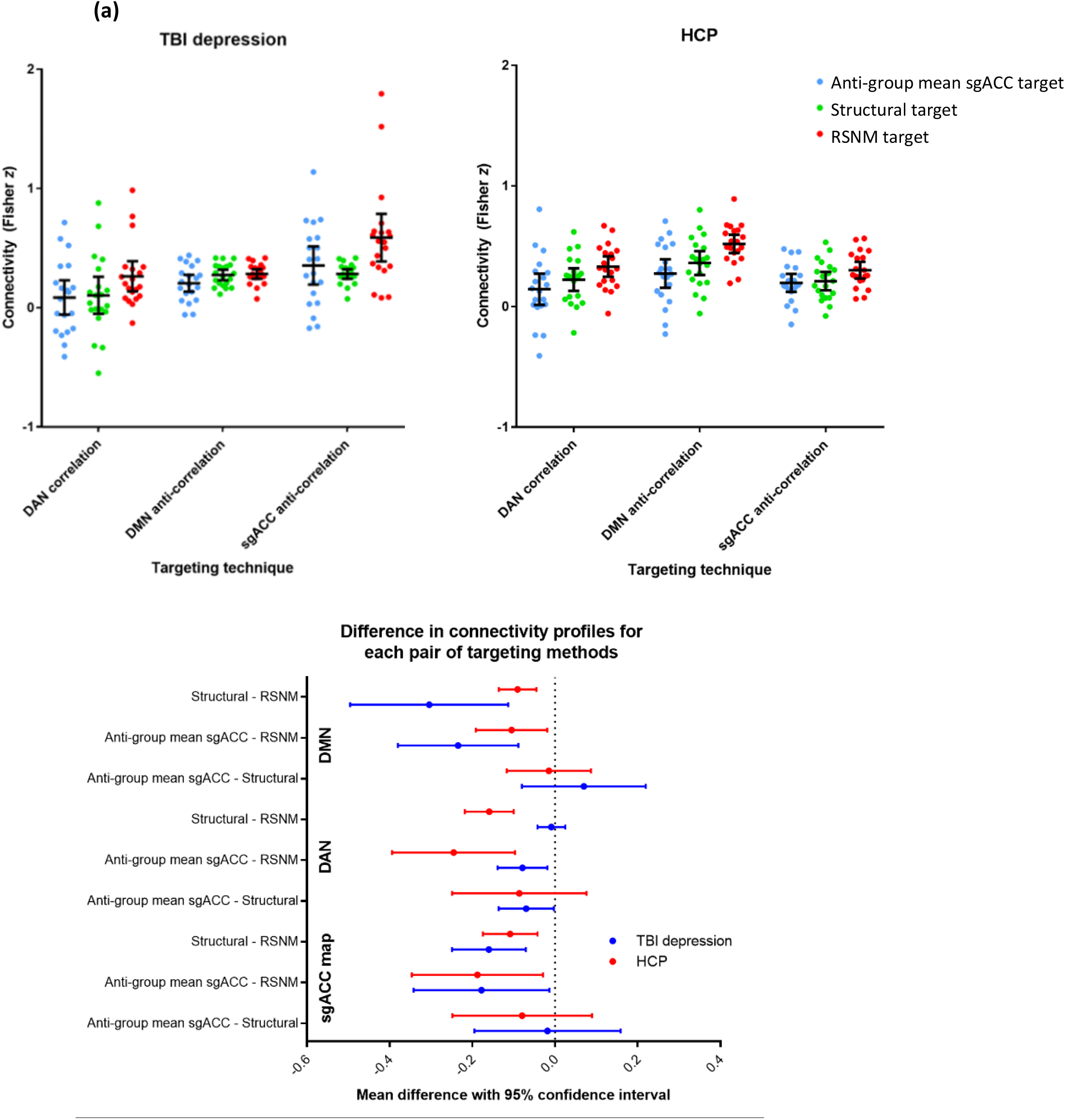
Functional connectivity of targets yielded by the three approaches. **(a)** Functional connectivity of RSNM, anti-group mean sgACC, and structural targets with DAN, DMN, and the normative sgACC connectivity map in each group. **(b)** Differences in connectivity profiles between the three potential targeting methods. On most metrics, RSNM targets showed significantly stronger connectivity with all three regions of interest.

### 3.3 Spatial distribution of derived targets

In both groups, RSNM-based target coordinates were spatially distinct from both comparator targets with 95% confidence intervals that were greater than zero (figure 4a-4b). The structural target was significantly closer to the RSNM target than to the anti-group mean sgACC target in both the TBI-D (p=0.006) and HCP (p=4×10^−5^) groups. The anti-group mean sgACC targets also showed wider variance between subjects than the RSNM targets for both the TBI-D (F=0.4, p=0.01) and HCP (F=0.4, p=0.03) groups (Table S2). The anatomical locations of targets generated by the different methods are depicted in figure 3d along with an example of approximate predicted stimulation volumes for each target in one representative subject. Thus, RSNM targets were less variable anatomically than anti-group mean sgACC targets.

**Figure 4:**
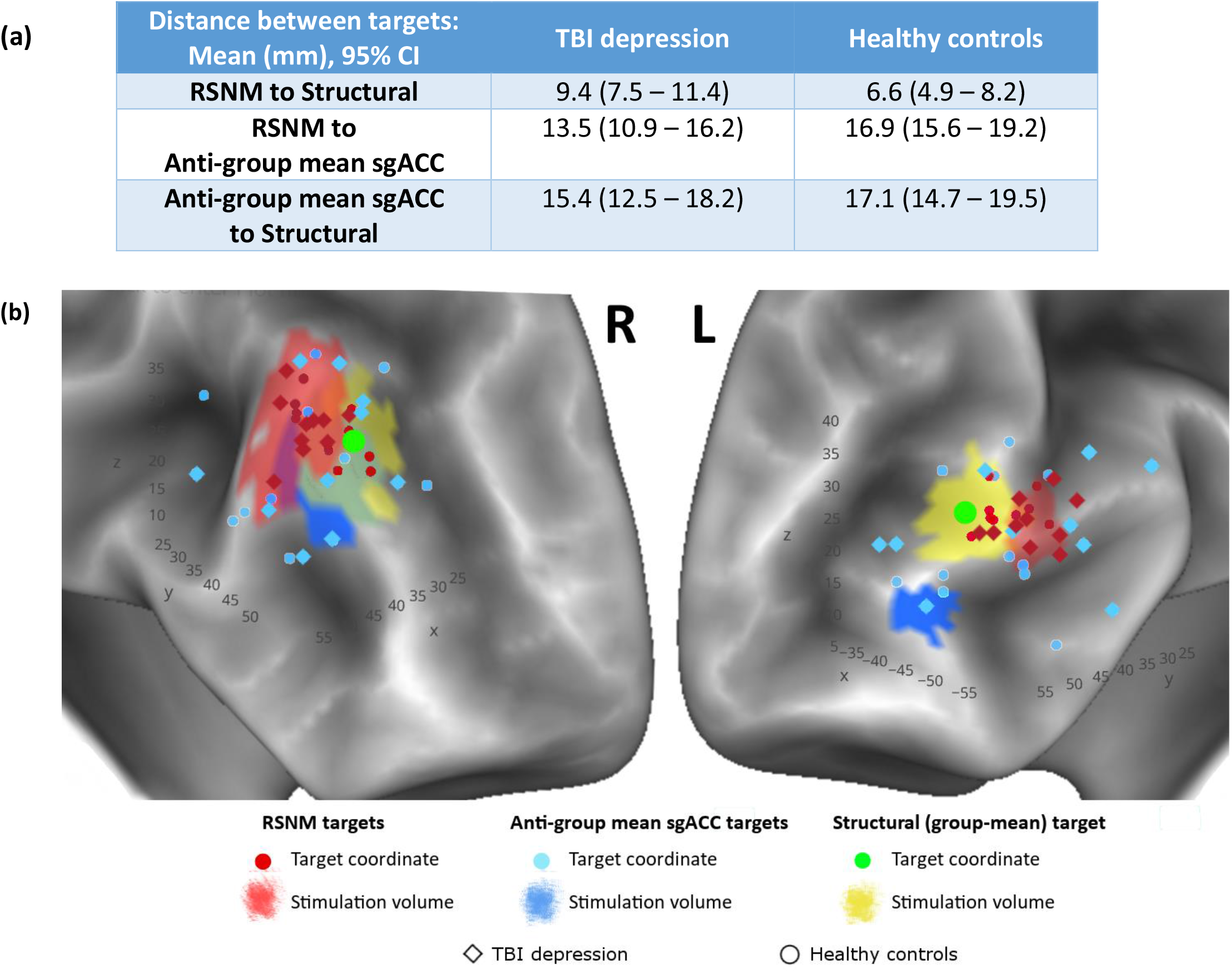
Anatomical distributions of targets yielded by the three approaches. **(a)** Mean and 95% confidence interval for the Euclidean distance between target coordinates generated by the different methods. **(b)** 3D scatter plot of the target sites generated by the different methods to illustrate approximate spatial distribution of targets. Background is a representative example of a single-subject surface reconstruction with approximate predicted stimulation volumes in that subject (in the TBI depression group). These approximate stimulation volumes are cortical surface projections of the estimated 15-mm sphere centered at the stimulation site for that subject; shapes are asymmetric and irregular due to normal variation in cortical surface anatomy. Across all subjects and in this representative example, RSNM-based actual stimulation sites were different from the structural group-mean site (green dot).

### 3.4 Stability of Connectivity and Target Location before vs. after RSNM targeted rTMS Treatment in TBI-D patients

13 TBI depression patients were scanned again after a full course of active rTMS (n=9) or sham rTMS (n=4). Connectivity with the normative sgACC seed map remained relatively stable for the RSNM target and the structural target (figure 6a). Anti-group mean sgACC targets, by contrast, showed significantly different connectivity profiles between pre-treatment and post-treatment scans (p=0.03). These results were unchanged when repeating the analysis after controlling for active versus sham stimulation (p=0.03), and there was no significant effect of treatment group (p=0.59).

Between the two scan sessions (pre- and post-treatment), the mean absolute Euclidean distance change in target coordinates was 6.6 mm for RSNM targets and 17.7 mm for anti-group mean sgACC targets (Wilcoxon matched-pairs signed rank test p<10^−4^, figure 6b). These results did not differ when repeating the analysis after controlling for active versus sham stimulation (p<10^−3^), and there was no significant effect of treatment group (p=0.14).

Thus, after active or sham rTMS at the RSNM target, the location of the RSNM targets remained more stable than the anti-group mean sgACC targets. Consistency of connectivity was similar between RSNM targets and structural targets. Of note, we did not directly assess stability of the location of the anti-group mean sgACC targets before and after stimulation at these targets because no stimulation was performed at anti-group mean sgACC targets.

### 3.5 Target engagement: treatment-induced change in connectivity

In comparison with sham, active rTMS was associated with large connectivity changes in several of the *a priori* ROI pairs, including DMN to sgACC, Left to Right stimulation site, and Left stimulation site to sgACC. There were also large changes in within-ROI connectivity in both stimulation sites and sgACC (Fig. 5c). Pre-treatment and post-treatment connectivity of each stimulation site with each of the 17 Yeo networks is depicted in Figure S2.

**Figure 5:**
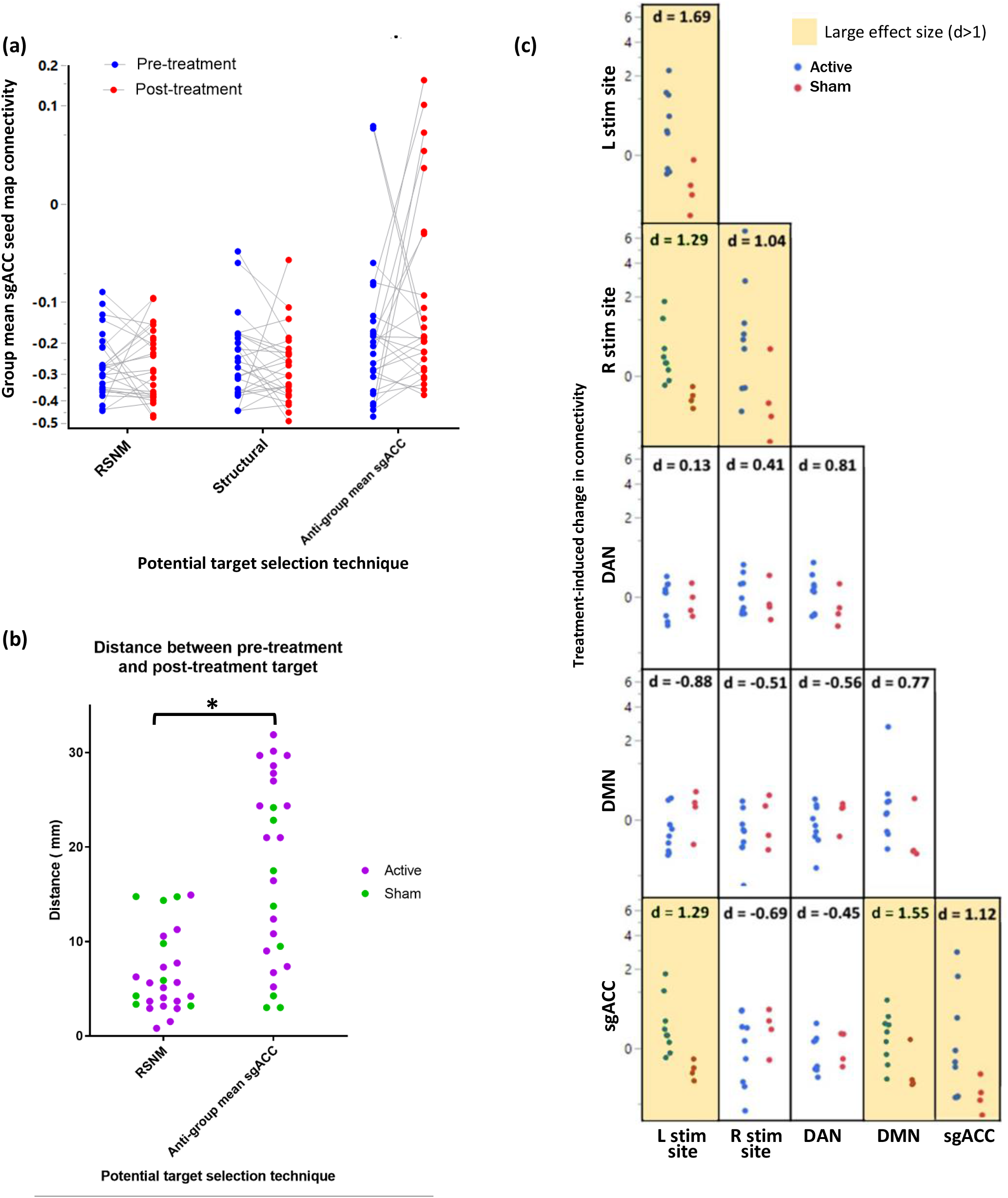
**(a) Change in connectivity profile of left- and right-sided potential targets identified based on pre-treatment and post-treatment scans**. Group mean sgACC seed map connectivity remained relatively stable for RSNM targets and structural targets but was more unstable for the potential anti-group mean sgACC targets. Each symbol represents 1 target; there were 2 targets per subject (right and left) x 13 subjects who underwent either active or sham rTMS treatment. **(b) Spatial change in target coordinates between pre-treatment and post-treatment scans**. After a course of RSNM-targeted treatment, the location of the potential anti-group mean sgACC target sites changed significantly more than the location of RSNM-based target sites (p < 0.0001, Wilcoxon matched-pairs signed-rank test). **(c) Treatment-induced change in connectivity within and between *a priori* ROIs**. Active treatment was associated with connectivity changes within and between stimulation sites, sgACC, DAN, and DMN. Magnitude of change is quantified using Cohen’s *d* (adapted with permission from Siddiqi *et al*, J Neurotrauma 2019).

Results and statistical methods for exploratory analyses are detailed in the supplement. First, treatment-induced connectivity change was compared between active and sham groups. Partial Spearman correlation was computed between group and post-treatment connectivity after controlling for pre-treatment connectivity (Figure S3). For the right stimulation site, there was a decrease in FC with the cingulo-opercular network parcel (rho=-0.56) and the parieto-occipital DAN parcel (rho= -0.55), increase in FC with the parahippocampal/retrosplenial DMN parcel (rho=0.65) (Figure S3a), and increase in FC with a voxel cluster in the left ventral hippocampus (r>0.7, corrected p<0.05) (Figure S3b). For the left stimulation site, active versus sham treatment led to a trend towards decreased parieto-occipital DAN connectivity and increased prefrontal/parietal DMN connectivity (Figure S3c), as well as a decrease in FC with a voxel cluster in the dorsomedial prefrontal cortex (r>0.7, corrected p<0.05) (Figure S3d).

### 3.6 Baseline predictors of clinical efficacy

For both stimulation sites, rTMS treatment efficacy was related to baseline connectivity of the stimulation site. For the right stimulation site, antidepressant response was significantly predicted by baseline anti-correlation with bilateral sgACC, anti-correlation with motor cortex, and positive correlation with dorsal ACC (corrected p<0.05) (figure 6a). For the left stimulation site, antidepressant response to rTMS was predicted by baseline correlation with right precuneus and anti-correlation with right sgACC, bilateral lateral parietal lobe, and bilateral dorsomedial prefrontal regions traditionally associated with the resting-state salience network (corrected p<0.05) (Figure 6b). Permutation testing confirmed that this whole-brain map was stronger than expected by chance for the right stimulation site (p = 0.04), but not the left stimulation site (p = 0.27).

**Figure 6:**
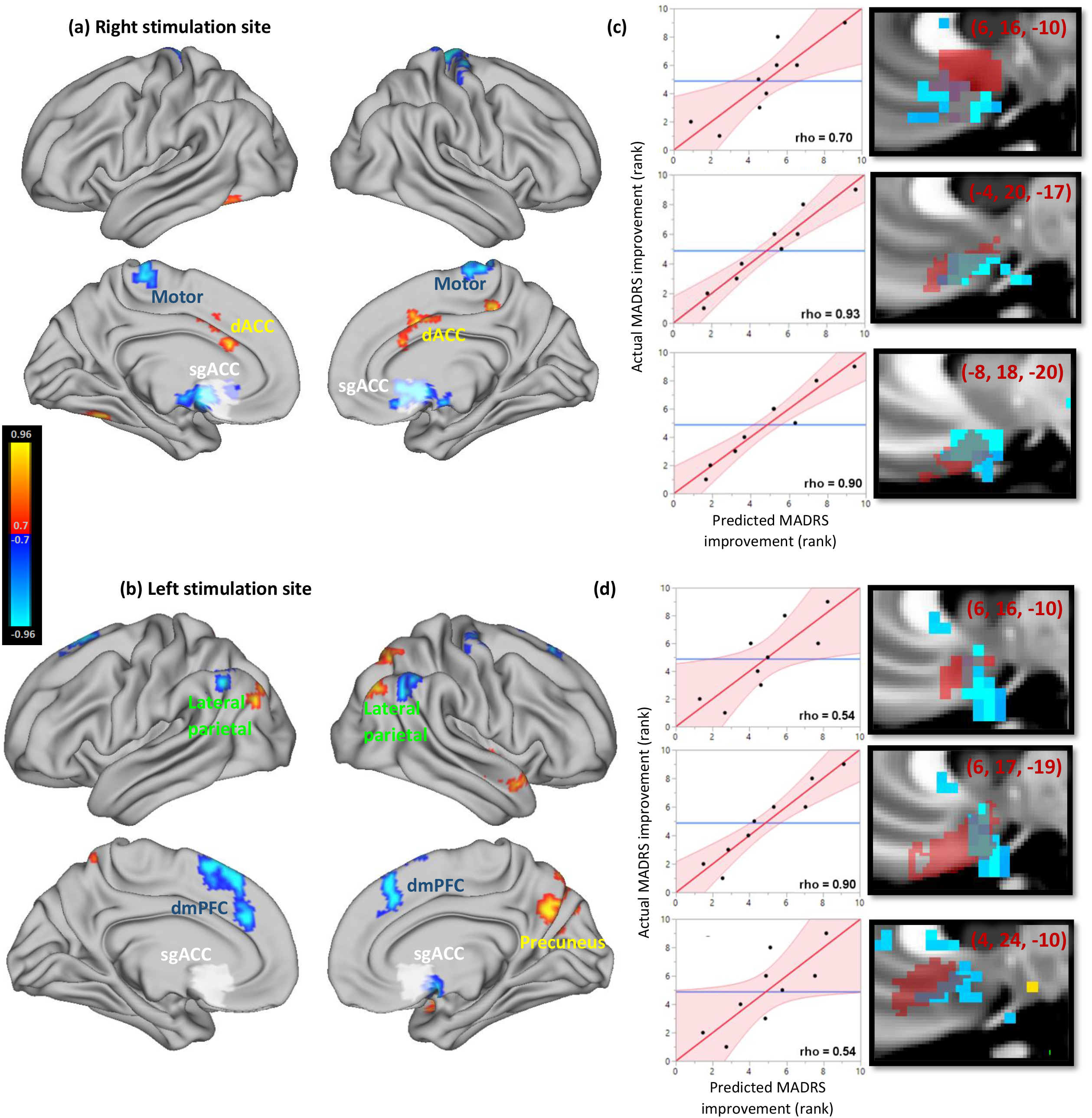
Connectivity profile associated with increased antidepressant efficacy of stimulation sites. White regions depict the *a priori* subgenual ROI. Clusters detected with threshold of r > 0.8 (uncorrected p < 0.001), minimum extent of 729 mm^3^, and cluster significance defined at p < 0.05. **(a)** Antidepressant response was significantly predicted by right stimulation site anti-correlation with bilateral sgACC, anti-correlation with motor cortex, and positive correlation with dorsal ACC (corrected p<0.05). **(b)** Antidepressant response was significantly predicted by left stimulation site correlation with right precuneus and anti-correlation with right sgACC, bilateral lateral parietal lobe, and bilateral dorsomedial prefrontal cortex (corrected p<0.05). **(c-d)** Antidepressant response was inversely related to seed-based connectivity of both stimulation sites with contralateral subgenual regions defined by a more recent cortical parcellation. Left panels depict the relationship between predicted and actual MADRS improvement, while right panels depict the overlap between the corresponding ROI (red) and the voxels whose stimulation site anti-correlation predicts MADRS improvement (blue). MNI coordinates of the center of each ROI are reported in maroon. **(c)** Right stimulation site connectivity with the *a priori* subgenual ROI was significantly predictive of antidepressant response (rho = 0.70, p = 0.035). This relationship was stronger when using exploratory subgenual ROIs based on a more recent cortical parcellation (rho = 0.90 and 0.93, p = 0.0009 and 0.0003). **(d)** Left stimulation site connectivity with the *a priori* subgenual ROI appeared to predict antidepressant response, but this relationship did not reach significance (rho = 0.54, p = 0.13). One of the two exploratory subgenual ROIs was significantly predictive of antidepressant response (rho = 0.90, p = 0.0009).

Antidepressant response was negatively correlated with the right-sided stimulation site’s FC with the *a priori* subgenual ROI (Figure 6c, top panel; Spearman rho = 0.70, p = 0.03). The left-sided stimulation site showed a trend in the same direction, but did not reach significance (figure 6d, top panel).

The voxel-wise maps of connections that predicted antidepressant response (Figures 6a and 6b) suggested that this effect was more prominent for specific subgenual regions that were only partially overlapping with the predefined sgACC ROI (Figures 6c and 6d, bottom two panels). To explore this further, the ROI-based analysis was repeated *post hoc* using recent sub-classified sgACC parcels^37^. Stimulation site connectivity with contralateral sgACC regions revealed strong predictive value for post-treatment MADRS (right stimulation site: rho = 0.90 and 0.93, p = 0.001 and 0.0003; left stimulation site: rho = 0.90 and 0.54, p = 0.001 and 0.13, respectively). Treatment efficacy was thus predictable using baseline stimulation site connectivity.

## 4. Discussion

Our findings suggest that individualized RSNM may be used to reliably identify rTMS targets based on their connectivity profile. We identified subject-specific rTMS targets at the networks that are likely being approximated by sgACC connectivity maps, which have previously been shown to predict efficacy of rTMS for major depression^5^. These target coordinates were stable and spatially distinct from prior approaches. Furthermore, these individualized RSNM-based targets showed stronger functional connectivity with the intended network targets than other candidate rTMS targets, even when these networks are defined conservatively based on consensus group-mean maps. Furthermore, the RSNM-based targets approximated the sgACC connectivity map more effectively than individualized targets generated using the previously-proposed anti-group mean sgACC approach. While it appears counter-intuitive that sgACC-based targets were less connected with a map generated using sgACC as a seed, this may be because the anti-group mean sgACC approach appears to generate unreliable targets. This is consistent with our hypothesis that our RSNM-based approach would identify a target that approximates the sgACC connectivity profile more effectively than a group-based sgACC seed.

Our proposed targeting approach was based on individualized mapping of DAN and DMN. The involvement of these networks in depression treatment may be related to dysfunctional interactions between externally-oriented attention-switching, which involves the DAN, and internally-oriented emotion engagement, which involves the DMN^38^. Such interactions appear to be affected in major depression^39^ and are modulated by deep brain stimulation of the sgACC^40^. This is consistent with our finding that antidepressant response was predicted by stimulation site connectivity with a large subgenual region. Treatment was also associated with changes in subgenual connectivity to itself, to the left DAN stimulation site, and to the DMN. This further suggests that our targeting approach may indirectly identify a network that modulates subgenual connectivity.

Nevertheless, our choice of DAN and anti-DMN targeting remains speculative in the absence of a head-to-head trial of antidepressant efficacy in comparison to rTMS applied to other targets. In addition, there are several existing approaches to individualized RSN mapping^6,16,17^ and we did not assess which approach best predicts neurophysiological and clinical response. There are also several approaches to resting-state fMRI pre-processing; for instance, our use of global signal regression may affect the identification of anti-correlated networks^41^. Similarly, there are several approaches to TMS-induced electric field modeling, but we chose not to use individualized finite element modeling because this method has not yet been validated for functional connectivity analyses. Careful validation of these techniques may help to further optimize our methods.

Our interpretation of treatment-induced changes are limited by small sample size. This may increase the risk of a false negative result due to lack of power or false positive results due to chance. Furthermore, we only assessed bilateral stimulation, not unilateral, and there may be complex interactions between the two stimulation sites, the significance of which is uncertain. This does not affect our reliability assessments, but does limit our ability to confirm whether the neurophysiological and clinical effects are consistent with our hypotheses. Importantly, prospective studies comparing unilateral stimulation vs. bilateral stimulation will be required to disentangle the neurophysiological effects of the bilateral stimulation employed in this study. It is not known whether the approach to selecting a left excitatory stimulation site should be the same as the approach used to selecting a right inhibitory stimulation site, since stimulation of the two hemispheres may have different effects^42^.

Further research will be required before these findings can be considered to be generalizable. This study was conducted using cutting-edge MRI scanners and recently-optimized scan protocols, so it remains unclear whether similar results can be achieved using more readily-available equipment. The patient population was also carefully selected as patients with relatively mild TBI and clear major depressive symptoms; its applicability to primary major depression or moderate/severe TBI requires further investigation.

Despite these limitations, our results support the emerging notion that variability in effects of rTMS may be related to inter-individual variability in functional topography of the DLPFC^1,43,44^. While the clinical implications of individualized RSN-based targeting are not yet clear, this method yields targets that are consistently connected to regions that have been implicated in antidepressant response to rTMS, including the sgACC. Stimulation of these targets also appears to modulate these key regions in a manner that is related to antidepressant response. This should help to inform an alternative and possibly more rational approach to prospective individualized target selection in future rTMS studies as well as retrospective analysis of results from existing studies.

In conclusion, the use of individualized RSN mapping for identification of distinct patient-specific rTMS targets may represent a promising method for reducing variability in targeting rTMS. This lays the foundation for development of more robust approaches for personalized medicine in neuromodulation.

## Acknowledgements

We would especially like to thank the experimental participants. We thank Dr. Abraham Snyder for logistical support with implementation of individualized RSN mapping. We thank Dr. Benjamin Srivastava for assisting with clinical assessments. We thank Linda Hood for technical support with MRI equipment. We thank Drs. Sindhu Jacob and Martin Wice for referring participants. We additionally acknowledge valuable discussions with Drs. Michael Fox, Steven Petersen, Charles Conway, Eric Wasserman, Sarah Lisanby, Bruce Luber, Andrew Drysdale, Irving Reti, Vani Rao, Maurizio Corbetta, Gordon Shulman, and Abraham Snyder which led to the conception and refinement of the study design.

## Author Contributions

SHS, DLB, ARC, and NTT designed the clinical protocol. ARC provided rTMS expertise and equipment as well as functional connectivity expertise. CDH provided expertise and novel analytical scripts for individualized RSN mapping and functional connectivity analysis. ECL provided expertise for clinical implementation of RSN mapping techniques. SK provided expertise and developed scripts for functional connectivity processing. SHS coordinated the study, recruited subjects, administered treatments, processed MRI data, conducted functional connectivity analyses, and conducted all statistical analyses. NTT, CDH, and XH also administered study treatments. SHS, NTT, LT, and PS acquired MRI scans and conducted clinical assessments. SHS, DLB, and ARC wrote the manuscript with intellectual contributions from all authors.

## Conflicts of interest

SHS serves as a clinical consultant for Kaizen Brain Center. SHS has received research support from Neuronetics Inc. The present work was not supported by any of these entities.

DLB has served as a consultant for Pfizer Inc, Intellectual Ventures, Signum Nutralogix, Kypha Inc, Sage Therapeutics, iPerian Inc, Navigant, Avid Radiopharmaceuticals (Eli Lilly & Co), the St Louis County Public Defender, the United States Attorney’s Office, the St Louis County Medical Examiner, GLG, Stemedica, and Luna Innovations. DLB holds equity in the company Inner Cosmos. DLB receives royalties from sales of Concussion Care Manual (Oxford University Press). No conflicts of interest with the presented work. DLB is an employee of the Department of Defense; the views expressed here do not reflect those of the Uniformed Services University of the Health Sciences, the US Department of Defense, or the US Government.

ECL holds equity in the companies Neurolutions and Inner Cosmos.

CDH, ECL, and SHS hold intellectual property related to the use of RSNM to target TMS.

The remaining authors report no conflicts of interest.

## References

1. Luber BM, Davis S, Bernhardt E, et al. Using neuroimaging to individualize TMS treatment for depression: Toward a new paradigm for imaging-guided intervention. Neuroimage. 2017;148:1– 7. doi.org/10.1016/j.neuroimage.2016.12.083 10.1016/j.neuroimage.2016.12.083. Epub 2017 Jan 3.

2. Fitzgerald PB, Hoy K, McQueen S, et al. A randomized trial of rTMS targeted with MRI based neuro-navigation in treatment-resistant depression. Neuropsychopharmacology. 2009;34(5):1255–1262. doi.org/10.1038/npp.2008.233

3. Fox MD, Buckner RL, Liu H, Chakravarty MM, Lozano AM, Pascual-Leone A. Resting-state networks link invasive and noninvasive brain stimulation across diverse psychiatric and neurological diseases. Proc Natl Acad Sci U S A. 2014;111(41):E4367–4375. doi.org/10.1073/pnas.1405003111

4. Fox MD, Buckner RL, White MP, Greicius MD, Pascual-Leone A. Efficacy of transcranial magnetic stimulation targets for depression is related to intrinsic functional connectivity with the subgenual cingulate. Biol Psychiatry. 2012;72(7):595–603. doi.org/10.1016/j.biopsych.2012.04.028 10.1016/j.biopsych.2012.04.028. Epub 2012 Jun 1.

5. Weigand A, Horn A, Caballero R, et al. Prospective Validation That Subgenual Connectivity Predicts Antidepressant Efficacy of Transcranial Magnetic Stimulation Sites. Biol Psychiatry. 2018;84(1):28–37. doi.org/10.1016/j.biopsych.2017.10.028

6. Cash RFH, Zalesky A, Thomson RH, Tian Y, Cocchi L, Fitzgerald PB. Subgenual Functional Connectivity Predicts Antidepressant Treatment Response to Transcranial Magnetic Stimulation: Independent Validation and Evaluation of Personalization. Biol Psychiatry. 2019;86(2):e5–e7. doi.org/10.1016/j.biopsych.2018.12.002

7. Cash RFH, Weigand A, Zalesky A, et al. Using Brain Imaging to Improve Spatial Targeting of Transcranial Magnetic Stimulation for Depression. Biol Psychiatry. 2020. doi.org/10.1016/j.biopsych.2020.05.033

8. Fox MD, Liu H, Pascual-Leone A. Identification of reproducible individualized targets for treatment of depression with TMS based on intrinsic connectivity. Neuroimage. 2013;66:151–160. doi.org/10.1016/j.neuroimage.2012.10.082

9. Williams NR, Sudheimer KD, Bentzley BS, et al. High-dose spaced theta-burst TMS as a rapid-acting antidepressant in highly refractory depression. Brain. 2018;141(3):e18. doi.org/10.1093/brain/awx379

10. Ning L, Makris N, Camprodon JA, Rathi Y. Limits and reproducibility of resting-state functional MRI definition of DLPFC targets for neuromodulation. Brain Stimul. 2019;12(1):129–138. doi.org/10.1016/j.brs.2018.10.004

11. Cash RFH, Cocchi L, Lv J, Fitzgerald PB, Zalesky A. Functional Magnetic Resonance Imaging–Guided Personalization of Transcranial Magnetic Stimulation Treatment for Depression. JAMA Psychiatry. 2020. doi.org/10.1001/jamapsychiatry.2020.3794

12. Mueller S, Wang D, Fox MD, et al. Reliability correction for functional connectivity: Theory and implementation. Hum Brain Mapp. 2015;36(11):4664–4680. doi.org/10.1002/hbm.22947

13. Laumann TO, Gordon EM, Adeyemo B, et al. Functional System and Areal Organization of a Highly Sampled Individual Human Brain. Neuron. 2015;87(3):657–670. doi.org/10.1016/j.neuron.2015.06.037

14. Glasser MF, Coalson TS, Robinson EC, et al. A multi-modal parcellation of human cerebral cortex. Nature. 2016;536(7615):171–178. doi.org/10.1038/nature18933

15. Hacker CD, Laumann TO, Szrama NP, et al. Resting state network estimation in individual subjects. Neuroimage. 2013;82:616–633. doi.org/10.1016/j.neuroimage.2013.05.108

16. Wang D, Buckner RL, Fox MD, et al. Parcellating cortical functional networks in individuals. Nat Neurosci. 2015;18(12):1853–1860. doi.org/10.1038/nn.4164

17. Birn RM, Molloy EK, Patriat R, et al. The effect of scan length on the reliability of resting-state fMRI connectivity estimates. Neuroimage. 2013;83:550–558. doi.org/10.1016/j.neuroimage.2013.05.099

18. Lee MH, Miller-Thomas MM, Benzinger TL, et al. Clinical Resting-state fMRI in the Preoperative Setting: Are We Ready for Prime Time? Top Magn Reson Imaging. 2016;25(1):11–18. doi.org/10.1097/RMR.0000000000000075

19. Siddiqi SH, Trapp NT, Hacker CD, et al. Repetitive Transcranial Magnetic Stimulation with Resting-State Network Targeting for Treatment-Resistant Depression in Traumatic Brain Injury: A Randomized, Controlled, Double-Blinded Pilot Study. J Neurotrauma. 2019;36(8):1361–1374. doi.org/10.1089/neu.2018.5889

20. Gordon EM, Laumann TO, Gilmore AW, et al. Precision Functional Mapping of Individual Human Brains. Neuron. 2017;95(4):791–807 e797. doi.org/10.1016/j.neuron.2017.07.011

21. Greicius MD, Flores BH, Menon V, et al. Resting-state functional connectivity in major depression: abnormally increased contributions from subgenual cingulate cortex and thalamus. Biol Psychiatry. 2007;62(5):429–437. doi.org/10.1016/j.biopsych.2006.09.020

22. Fox MD, Snyder AZ, Vincent JL, Corbetta M, Van Essen DC, Raichle ME. The human brain is intrinsically organized into dynamic, anticorrelated functional networks. Proc Natl Acad Sci U S A. 2005;102(27):9673–9678. doi.org/10.1073/pnas.0504136102

23. Chai XJ, Castanon AN, Ongur D, Whitfield-Gabrieli S. Anticorrelations in resting state networks without global signal regression. Neuroimage. 2012;59(2):1420–1428. doi.org/10.1016/j.neuroimage.2011.08.048

24. Wong CW, Olafsson V, Tal O, Liu TT. Anti-correlated networks, global signal regression, and the effects of caffeine in resting-state functional MRI. Neuroimage. 2012;63(1):356–364. doi.org/10.1016/j.neuroimage.2012.06.035

25. Gordon EM, Laumann TO, Adeyemo B, et al. Individual-specific features of brain systems identified with resting state functional correlations. Neuroimage. 2017;146(918-939):918–939. doi.org/10.1016/j.neuroimage.2016.08.032

26. Gordon EM, Laumann TO, Adeyemo B, Petersen SE. Individual Variability of the System-Level Organization of the Human Brain. Cereb Cortex. 2017;27(1):386–399. doi.org/10.1093/cercor/bhv239

27. Han K, Mac Donald CL, Johnson AM, et al. Disrupted modular organization of resting-state cortical functional connectivity in U.S. military personnel following concussive ‘mild’ blast-related traumatic brain injury. Neuroimage. 2014;84:76–96. doi.org/10.1016/j.neuroimage.2013.08.017 10.1016/j.neuroimage.2013.08.017. Epub 2013 Aug 20. From Duplicate 2 (Disrupted modular organization of resting-state cortical functional connectivity in U.S. military personnel following concussive ‘mild’ blast-related traumatic brain injury – Han, K; Mac Donald C L; Johnson, A M; Barnes, Y; Wierzechowski, L; Zonies, D; Oh, J; Flaherty, S; Fang, R; Raichle, M E; Brody D L) Han, Kihwan Mac Donald, Christine L Johnson, Ann M Barnes, Yolanda Wierzechowski, Linda Zonies, David Oh, John Flaherty, Stephen Fang, Raymond Raichle, Marcus E Brody, David L F32 NS062529/NS/NINDS NIH HHS/ S10 RR022984/RR/NCRR NIH HHS/ 1S10RR022984-01A1/RR/NCRR NIH HHS/ Neuroimage. 2014 Jan 1;84:76–96. doi:10.1016/j.neuroimage.2013.08.017. Epub 2013 Aug 20.

28. Han K, Chapman SB, Krawczyk DC. Disrupted Intrinsic Connectivity among Default, Dorsal Attention, and Frontoparietal Control Networks in Individuals with Chronic Traumatic Brain Injury. J Int Neuropsychol Soc. 2016;22(2):263–279. doi.org/10.1017/S1355617715001393

29. van der Horn HJ, Liemburg EJ, Scheenen ME, de Koning ME, Spikman JM, van der Naalt J. Graph Analysis of Functional Brain Networks in Patients with Mild Traumatic Brain Injury. PLoS One. 2017;12(1):e0171031. doi.org/10.1371/journal.pone.0171031

30. Caeyenberghs K, Verhelst H, Clemente A, Wilson PH. Mapping the functional connectome in traumatic brain injury: What can graph metrics tell us? Neuroimage. 2017;160:113–123. doi.org/10.1016/j.neuroimage.2016.12.003

31. Siddiqi SH. rTMS for major depression associated with TBI. 2016; osf.io/vjddq.

32. Van Essen DC, Ugurbil K, Auerbach E, et al. The Human Connectome Project: a data acquisition perspective. Neuroimage. 2012;62(4):2222–2231. doi.org/10.1016/j.neuroimage.2012.02.018

33. Power JD, Mitra A, Laumann TO, Snyder AZ, Schlaggar BL, Petersen SE. Methods to detect, characterize, and remove motion artifact in resting state fMRI. Neuroimage. 2014;84:320–341. doi.org/10.1016/j.neuroimage.2013.08.048

34. Siddiqi SH, Trapp NT, Shahim P, et al. Individualized Connectome-Targeted Transcranial Magnetic Stimulation for Neuropsychiatric Sequelae of Repetitive Traumatic Brain Injury in a Retired NFL Player. J Neuropsychiatry Clin Neurosci. 2019;31(3):254–263. doi.org/10.1176/appi.neuropsych.18100230

35. Blumberger DM, Maller JJ, Thomson L, et al. Unilateral and bilateral MRI-targeted repetitive transcranial magnetic stimulation for treatment-resistant depression: a randomized controlled study. J Psychiatry Neurosci. 2016;41(4):E58–66. doi.org/10.1503/jpn.150265

36. Yeo BT, Krienen FM, Sepulcre J, et al. The organization of the human cerebral cortex estimated by intrinsic functional connectivity. J Neurophysiol. 2011;106(3):1125–1165. doi.org/10.1152/jn.00338.2011 10.1152/jn.00338.2011. Epub 2011 Jun 8.

37. Schaefer A, Kong R, Gordon EM, et al. Local-Global Parcellation of the Human Cerebral Cortex from Intrinsic Functional Connectivity MRI. Cereb Cortex. 2018;28(9):3095–3114. doi.org/10.1093/cercor/bhx179

38. Hamilton JP, Furman DJ, Chang C, Thomason ME, Dennis E, Gotlib IH. Default-mode and task-positive network activity in major depressive disorder: implications for adaptive and maladaptive rumination. Biol Psychiatry. 2011;70(4):327–333. doi.org/10.1016/j.biopsych.2011.02.003

39. Kaiser RH, Andrews-Hanna JR, Wager TD, Pizzagalli DA. Large-Scale Network Dysfunction in Major Depressive Disorder: A Meta-analysis of Resting-State Functional Connectivity. JAMA Psychiatry. 2015;72(6):603–611. doi.org/10.1001/jamapsychiatry.2015.0071

40. Choi KS, Riva-Posse P, Gross RE, Mayberg HS. Mapping the “Depression Switch” During Intraoperative Testing of Subcallosal Cingulate Deep Brain Stimulation. JAMA Neurol. 2015;72(11):1252–1260. doi.org/10.1001/jamaneurol.2015.2564

41. Murphy K, Fox MD. Towards a consensus regarding global signal regression for resting state functional connectivity MRI. Neuroimage. 2017;154:169–173. doi.org/10.1016/j.neuroimage.2016.11.052

42. Chen J, Zhou C, Wu B, et al. Left versus right repetitive transcranial magnetic stimulation in treating major depression: a meta-analysis of randomised controlled trials. Psychiatry Res. 2013;210(3):1260–1264. doi.org/10.1016/j.psychres.2013.09.007

43. Sale MV, Mattingley JB, Zalesky A, Cocchi L. Imaging human brain networks to improve the clinical efficacy of non-invasive brain stimulation. Neurosci Biobehav Rev. 2015;57:187–198. doi.org/10.1016/j.neubiorev.2015.09.010

44. Opitz A, Fox MD, Craddock RC, Colcombe S, Milham MP. An integrated framework for targeting functional networks via transcranial magnetic stimulation. Neuroimage. 2016;127:86–96. doi.org/10.1016/j.neuroimage.2015.11.040

